# Optimizing Pedigrees: Using a Biasing System to Determine Likely Inheritance Systems

**DOI:** 10.1101/116160

**Authors:** Justin Ang

**Affiliations:** University of California, San Diego

## Abstract

Pedigrees, though straightforward and versatile, lack the ability to tell us information about many individuals. Though numerical systems have been developed, there is currently no system to quantify the probability of a pedigree following certain inheritance systems. My system intends to fulfill that chasm by creating a flexible numerical system and testing it for variance. First, my system attempts to adapt inheritance system to known pedigree data. Then, it calculates the difference between the calculated values and the known pedigree data. It aggregates these values, then it uses a chi-squared analysis in order to determine the likelihood of said inheritance system. This is done for many different systems, until we have a general idea of which systems are probable and which are not.

## Introduction

Pedigrees are fundamentally used to represent phenotypic data in a binary system. By using ’Black’ as affected and ‘White’ as unaffected, the pedigree system is able to cover a wide range of inheritance systems. It also gives a general idea of the dispersion of the phenotype in question. A simple improvement to this existing model would be to assign scores instead of coloring. These scores would keep track of the chance that individuals hold certain traits which can be used to easily calculate the probability of child generations. This approach has many limitations, but the idea of such an implementation is very interesting and applicable [1](Weighted-Based Statistics) (*T*^2^ Test)[2] (*M*_*QLS*_ Test)[3]. However, using scores can also be used to determine the chance of basic inheritance systems, which I will prove. By finding the variance between the expected value and observed values of models, a chi-squared analysis can be performed to obtain a p-value. This p-value will then allow us to determine if a system is probable or not.

My system and framework is very similar to other recent models. For example, a Bayesian framework[7] has been introduced to quantify pedigrees as well. My algorithm is not intended to compete with, nor beat those models. Alternatively, my model focuses on being able to quantify inheritance patterns. Of course, my model has many similarities to other work in the field due to the need for a numerical ordering system. As such, it carries many of the benefits and drawbacks that other numerical systems hold. I understand that others have tried to maximize positives and minimize negatives by introducing more layers of statistical analysis. The model I created was less rigorous in order to prove the benefits of determining inheritance patterns more easily. However, additional levels of statistics are definitely applicable to these models, and may be a point of interest for some.

## Methods

### Assumptions

This model depends on several assumptions:

1. We are only looking at a two-allele, one gene system that is governed by Mendelian inheritance.
  - This model assumes that for every subsequent level the genotypes of the parents is diluted by exactly 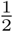. If the alleles are not separated, then there is no guarantee that these models will hold true. [7]
2. We are only looking at: Autosomal Recessive, Autosomal Dominant, X-linked Recessive, X-linked Dominant, and Y-linked inheritance patterns.
3. There are no abnormal females and males. The individuals have normal karyotypes. (XXY, XO, XXX, etc. are not present)
4. There is **full** penetrance.

### Basic Structure

Here, I will document key aspects of my representations. These are present in every implementation:

Each individual will be assigned **one** number. This number is responsible for representing the probability of affected/unaffectedness. This number will be also used to calculate offspring.

#### Signed numbers

Positive (0-2) = Affected Bias (Possibly Affected)

Very Positive (1-2) = Definitely Affected

Neutral (0) = No Bias

Negative (-1-0) = Unaffected Bias (Unaffected)

Bias represents the chance of being affected. Bias will be integral in calculating the probabilities of subsequent generations inheriting alleles.

##### Why is the Bias unbalanced?

The bias is unbalanced because Dominant and Recessive inheritance schemes are unbalanced. Giving the same weighting system, say, {-2, 2} fails to give us a distinction between the two. By adding favoritism to the bias of Dominant traits, we ensure that the variance between Recessive and Dominant models is significant.

#### Level Calculations

The biases of the parents will be summed and divided by two for every subsequent generation unless otherwise specified (X-Linked traits and Y-Linked traits). This is due to each parent only having a 50% chance of passing on their genotype under the aforementioned environment. This is essentially a Kinship Coefficient [1] calculation restricted to parents and children. *For every child, assign the biases based on the kinship Coefficient, where Ψ is the sum of the two parents’ biases.*

Essentially:

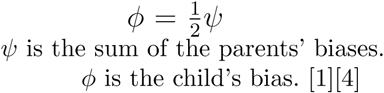

#### Self-Correcting

The algorithm will always prioritize data over calculations. Suppose a calculation for an individual gives us a value that is inconsistent with the known traits of the individual. We will use values based on the trait of the individual instead of the one we have calculated. This will only occur when we are mating an individual based on phenotype, and are unsure if they are homozygous or heterozygous. This is could be seen as following the same logic as the Collapsing Method Fundamental[1], albeit much simpler.

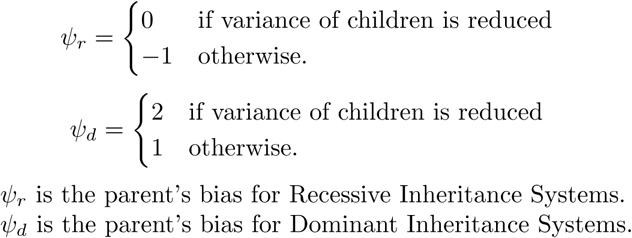

Repeat for all unsure individuals.

Since I use similar concepts behind my numerical system as many other studies, I will keep this section brief. I have attached a supplement for further reading into the rationale behind my methods. However, the specifics of my numerical analysis are not the core of my argument.

##### Autosomal Recessive

Assign a bias factor of +1 if the individual is affected. Assign a bias factor of -1 if the individual is definitely unaffected or unknown. Assign a bias factor of 0 if the individual is a carrier. If the I generation consists of two unaffected individuals, assign them both 0 unless otherwise specified. If two unaffected individuals produce an unaffected individual, assign the individual a bias of -0.33. If two unaffected individuals produce an affected individual, assign the individual a bias of +1 and assign the parents to have a bias of 0. For every child, assign the biases based on the kinship Coefficient, where *Ψ* is the sum of the two parents’ biases.

##### Autosomal Dominant

Assign a bias factor of +2 if the individual is fully affected. Assign a bias factor of 0 if the individual is unaffected. Assign a bias factor of +1 if the individual is unknown or a Heterozygote. If the I generation consists of affected individuals, assign them all +1 unless otherwise specified. If two unaffected individuals produce an unaffected individual, assign the individual a bias of -1. If the individuals produce an affected individual, assign the individual a bias of +1.33.

##### X-Linked Recessive

Assign a bias factor of +1 if the individual is affected. Assign a bias factor of 0 if the individual is a **female** carrier. Assign a bias factor of -1 if the individual is unaffected or unknown. **Males will have their mother’s bias**, instead of calculating the Kinship Coefficient. If the I generation consists of affected individuals, assign them all 1 unless otherwise specified. If two carrier individuals produce an unaffected **female**, assign the female a bias of -0.5. If the individuals produce an affected female, assign the individual a bias of +1.

##### X-Linked Dominant

Assign a bias factor of +2 if the individual is fully affected. Assign a bias factor of 1 if the individual is a **female** heterozygote. Assign a bias factor of 0 if the individual is unaffected. **Males will have their mother’s bias**, instead of calculating the Kinship Coefficient. If the I generation consists of affected individuals, assign them all +1 unless otherwise specified. If two heterozygous individuals produce an unaffected **female**, assign the female a bias of 0. If the individuals produce an affected female, assign the individual a bias of +1.5.

##### Y-Linked

Assign a bias factor of +2 if the individual is male and affected. In the impossible case where a female is affected or a carrier, assign it a bias of *∞*. Assign a bias factor of 0 to unaffected males, and all other females. **Males will have their father’s bias**. None of the individuals will follow the Kinship Coefficient model in this inheritance scheme.

### Using Models to Determine Inheritance System

The aforementioned models can be used to determine the inheritance type of an unknown pedigree. The method for doing so is as follows:

1. Attempt to fit each model to the given tree. Most models will not fit. In that case find the model that is the **closest** to the given tree. Closeness is defined as minimizing the difference between the calculated bias and the observed bias.
2. Test the sum of the bias of individuals in a set of parents minus the expected bias as calculated from the set of parents.
3. Sum the absolute value of these differences for every set of parents after the I generation. This is considered one system.
4. If we happen to cross a set of parents where the expected bias of the parents conflicts with the one we observe, assign the observed bias instead of the calculated one.
5. Square this value. This is our variance for the parent-child system.
6. Test these variance values against a chi-square analysis. Our Degrees of Freedom will be the number of parent-child systems minus 1.
7. If we are given information about heterozygotes, we assume unaffected and affected individuals to be homozygotes. However, phenotypes are prioritized. For example, carriers will not be heterozygous in a Autosomal Dominant scheme.

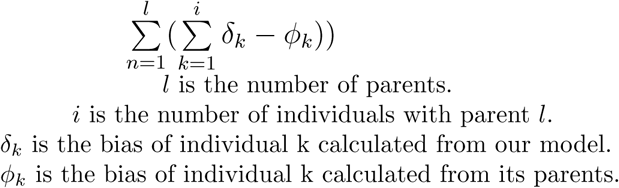

#### Null hypothesis

There is no significant difference between the data observed and the data expected. Therefore the proposed model is accurate with respect to our data. Assume p = 0.05 [5] for us to be able to reject the null hypothesis.

The reasoning for this system is explained in the “Further Explanation of Variance”, on page 13. I will now proceed to analyze a few pedigrees. For the sake of simplicity, the models will not have shading.

### Further Explanation of Variance

The variance calculation is as follows:

1. Generate models for all 5 inheritance systems.
2. Find the difference between expected value and observed value for children.
3. Sum all the child differences from a set of parents and square this value.
4. Compare to p-value. Degrees of freedom is the number of parent-child systems minus one.
5. If we are given information about heterozygotes, we assume unaffected and affected individuals to be homozygotes. Phenotypes are considered before heterozygosity.

#### Explanation

1. We must generate models to obtain our variance. Different models will create different expected values which will in turn create different variance.

2. In order to find our variance, we need to find the bias we expect minus the bias we are getting.

3. The variance is based on **parent sets**. Our calculation from two parents will give us the expected bias of all the children. In order to find the variance of a parent set, we must sum the individual variances of the children. If we only take the variance of one child without accounting for the other children, the variance will not be accurate.

We square the sums of the differences because we are treating a parent-child system collectively and are not looking at individuals.

4. The range of factors (individuals) increases as we increase our parent-child systems. Therefore, our Degrees of freedom will rise as we look at more systems.

5. If we know all the heterozygotes, we do not need to self-correct. Phenotypes are prioritized because heterozygotes take on different traits in different systems. If phenotypes for heterozygotes are ignored, then Dominant and Recessive schemes will have the same variance.

## Results

### Example 1 (Autosomal Recessive)

**Figure 1:**
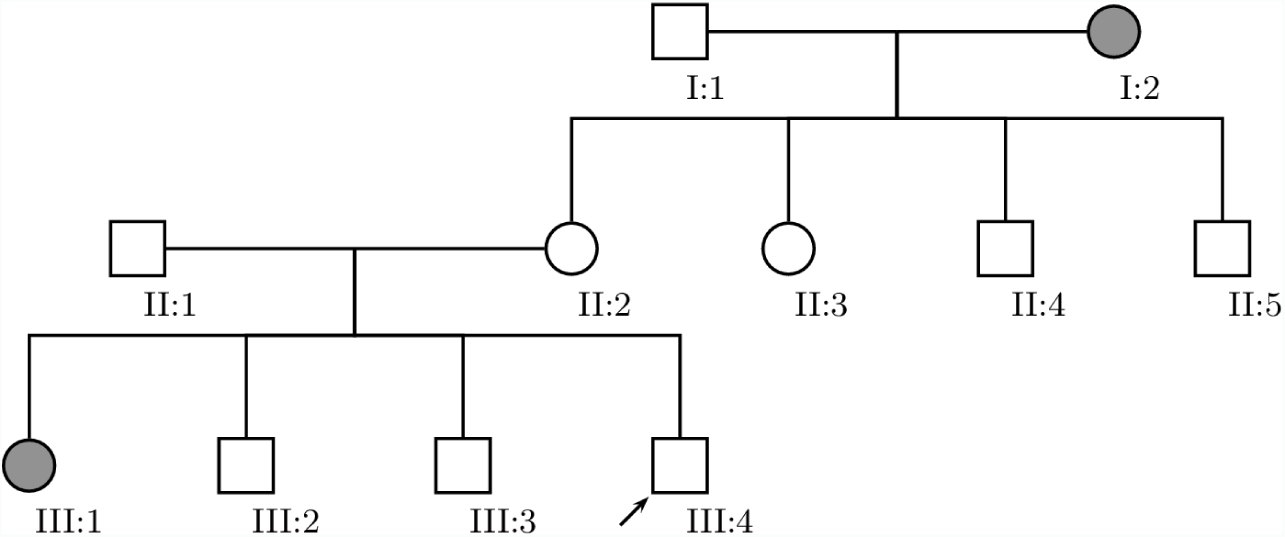
Example family that follows an Autosomal Recessive inheritance scheme. Generation II is unaffected, which is characteristic of recessive inheritance schemes.

**Figure 2:**
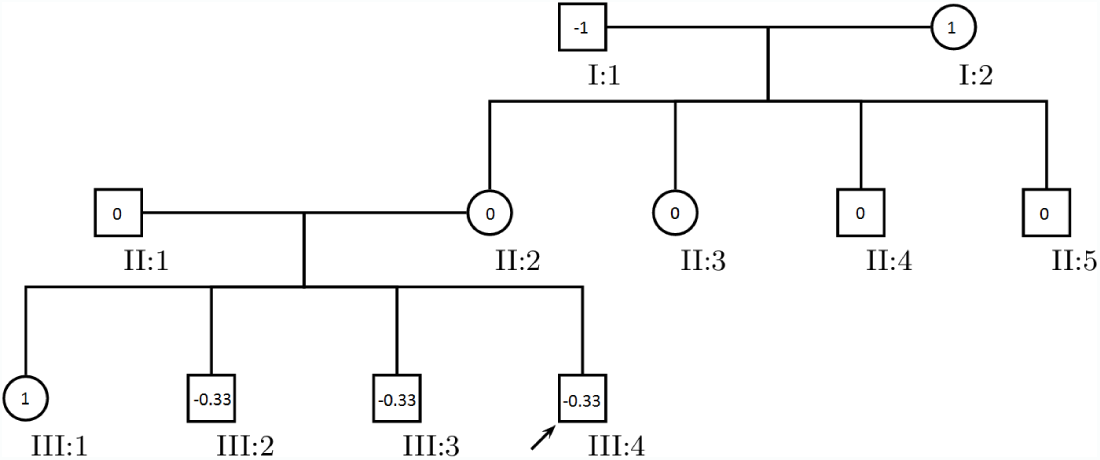
Assignment of bias to an Autosomal Recessive inheritance scheme.

Expected Bias from I Parents to II children: 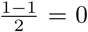

Bias from II children (II-2, II-3, II-4, II-5): (0-0) + (0-0) + (0-0) + (0-0) = 0^2^ = 0

Expected Bias from (II-1, II-2) Parents to III children: 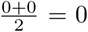

Bias from III children (III-1, III-2, III-3, III-4): (1-0) + (-0.33-0) + (-0.33-0) + (-0.33-0) = 0^2^ = 0

Total Bias from all levels: *|*0*|* + *|*0*|* = 0

**Figure 3:**
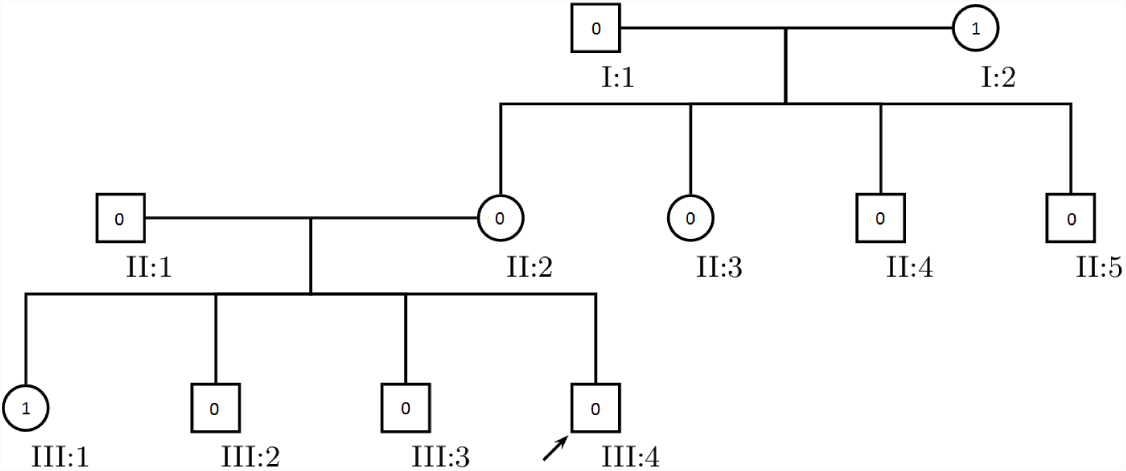
Assignment of bias to an Autosomal Dominant inheritance scheme.

Expected Bias from I Parents to II children: 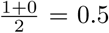

Bias from II children (II-2, II-3, II-4, II-5): (0-0.5) + (0-0.5) + (0-0.5) + (0-0.5) = −2^2^ = 4

Expected Bias from (II-1, II-2) Parents to III children: 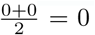

Bias from III children (III-1, III-2, III-3, III-4): (1-0) + (0-0) + (0-0) + (0-0) = 1^2^ = 1

Total Bias from all levels: 4 + 1 = 5

**Figure 4:**
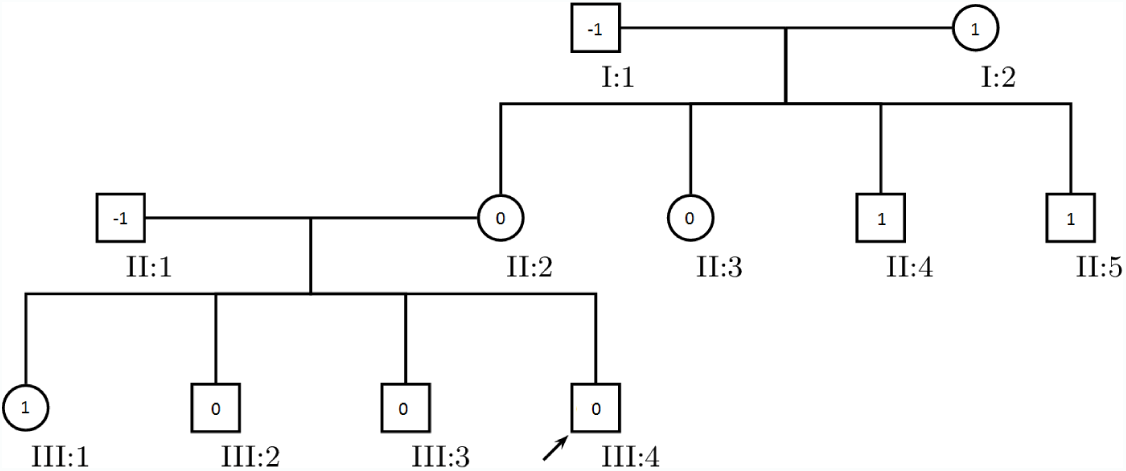
Assignment of bias to an X-Linked Recessive inheritance scheme.

Expected Bias from I Parents to II children: ♂ = 1, 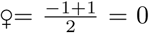

Bias from II children (II-2, II-3, II-4, II-5): (0-0) + (0-0) + (1-1) + (1-1) = 0

Expected Bias from (II-1, II-2) Parents to III children: ♂ = 0, 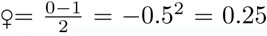

Bias from III children (III-1, III-2, III-3, III-4): (1-(-0.5)) + (0-0) + (0-0) + (0-0) = 1.5^2^ = 2.25

Total Bias from all levels: 0.25 + 2.25 = 2.50

**Figure 5:**
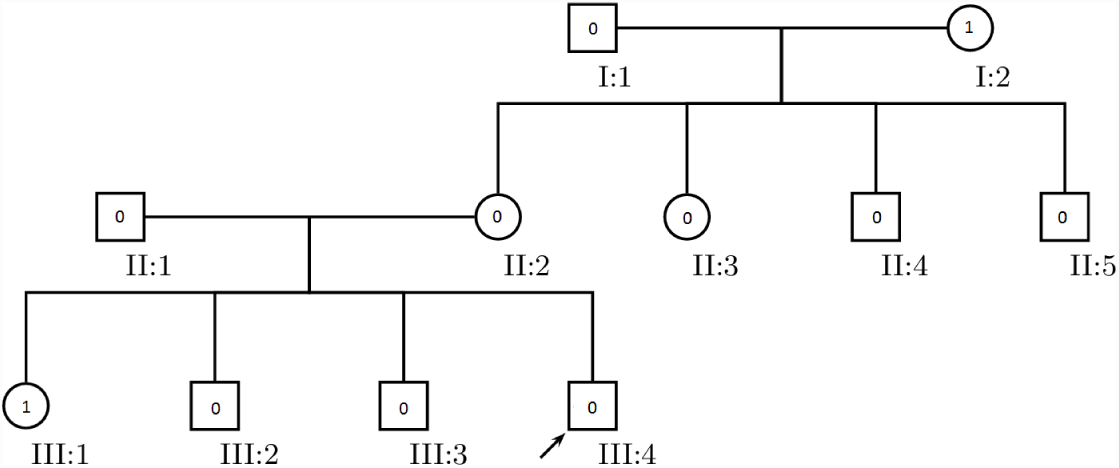
Assignment of bias to an X-Linked Dominant inheritance scheme.

Expected Bias from I Parents to II children: ♂ = 1, 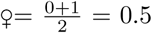

Bias from II children (II-2, II-3, II-4, II-5): (0-0.5) + (0-0.5) + (0-1) + (0-1) = −3^2^ = 9

Expected Bias from (II-1, II-2) Parents to III children: ♂ = 0, 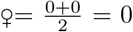

Bias from III children (III-1, III-2, III-3, III-4): (1-0) + (0-0) + (0-0) + (0-0) = 1^2^ = 1

Total Bias from all levels: 9 + 1 = 10

**Figure 6:**
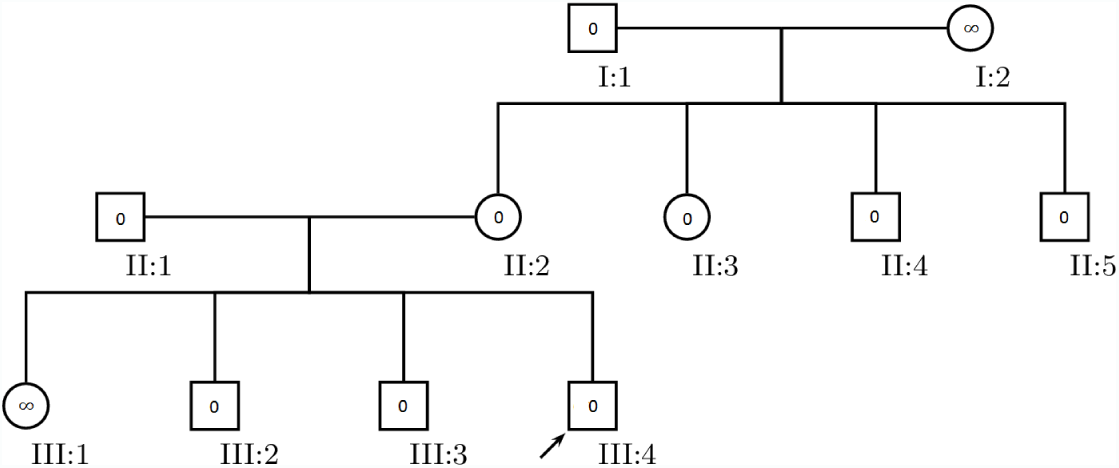
Assignment of bias to an Y-Linked inheritance scheme.

Expected Bias from I Parents to II children: ♂= 0, ♀= 0

Bias from II children (II-2, II-3, II-4, II-5): (0-0) + (0-0) + (0-0) + (0-0) = 0

Expected Bias from (II-s1, II-2) Parents to III children: ♂= 0, 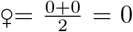

Bias from III children (III-1, III-2, III-3, III-4): (*∞*-0) + (0-0) + (0-0) + (0-0) = *∞*^2^= *∞*

Total Bias from all levels: 0 + *∞* = *∞*

After our calculations, we have the corresponding values for Autosomal Recessive, Autosomal Dominant, X-Linked Recessive, X-Linked Dominant, and Y-Linked: 0, 12, 9, 16, *∞*. Now, let us apply these values to a Chi-Square Distribution. Since we examined 2 parent-child systems (I-1/I-2 & II-1/II-2), our degrees of freedom is 2-1 = 1.

**Table 1:** Analysis of Inheritance Patterns for Example 1.

| Inheritance Pattern | $X^2$ Value | p-value | Probable? |
| --- | --- | --- | --- |
| Autosomal Recessive | 0 | 1 | yes |
| Autosomal Dominant | 5 | 0.025 | no |
| X-Linked Recessive | 2.50 | 0.114 | yes |
| X-Linked Dominant | 10 | 0.001 | no |
| Y-Linked | ∞ | 0 | no |

From this distribution, we can see that Autosomal Recessive and X-Linked Recessive are models that fail to reject the null hypothesis. However, Autosomal Recessive has a much higher confidence when applied to this pedigree. X-Linked Recessive should, in theory, be impossible. An affected female (III-2) cannot be born from an unaffected male (II-1) and unaffected female (II-2), as the female will always get a working allele from their father. However, this model accounts for error in results. Since this system is not meant to give a binary answer to inheritance systems, some inheritance systems may remain probable but extremely unlikely. Our rejection of the X-Linked Dominant inheritance scheme and Autosomal Dominant Scheme matches the pedigree. It is impossible for an affected female (III-1) to be born from an unaffected father (II-1). Additionally, Y-linked traits cannot be expressed by females, yet affected females are present in this pedigree. Therefore, this pedigree does not follow Y-Linked inheritance. My chi-squared test confirms this.

Another reason why impossible systems are still included is due to the small sample size. If we were to have more individuals, then our system will deduce with more certainty if a model fits. [5]

### Example 2 (Y-Linked, Chance of Autosomal)

Now, I will use a simpler example to prove that this method is accurate. Assume we know nothing but the phenotypes.

**Figure 7:**
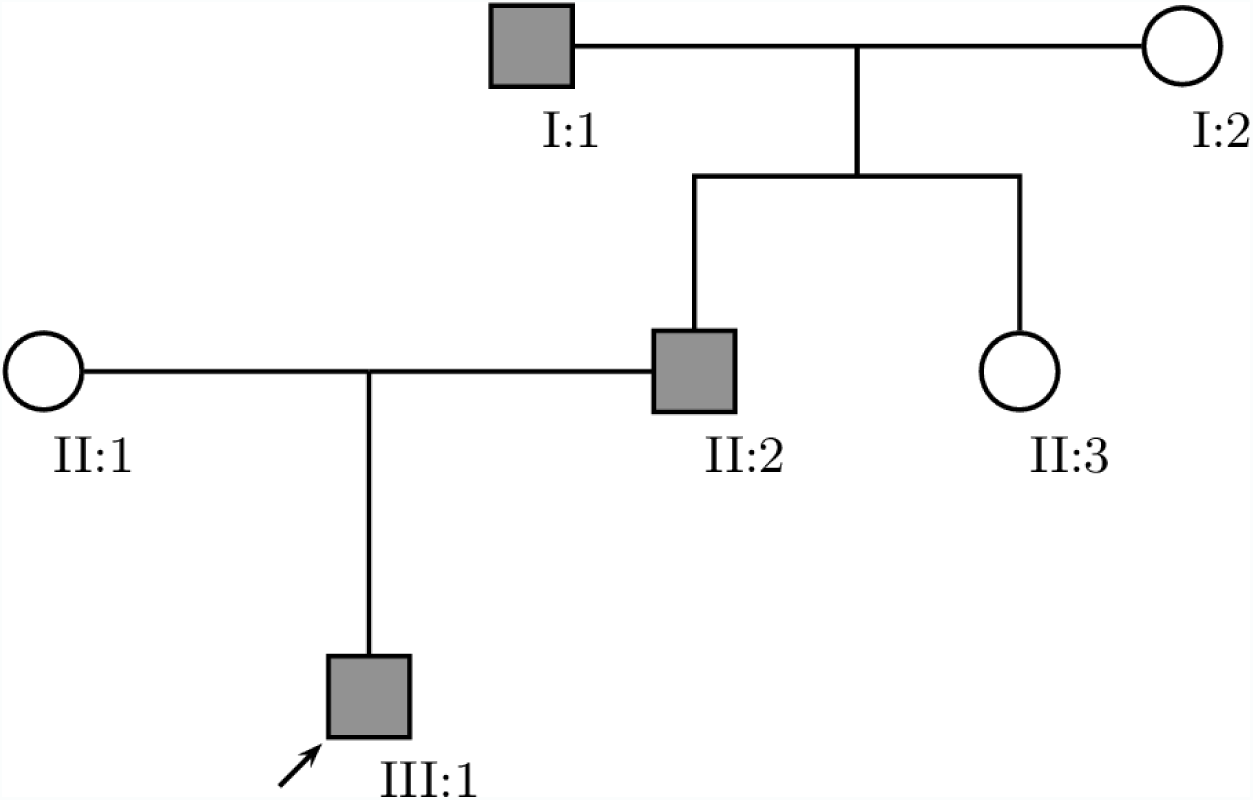
Example family that follows an Y-Linked inheritance scheme. Only males are affected in this pedigree, therefore Y-Linked is highly probable.

**Figure 8:**
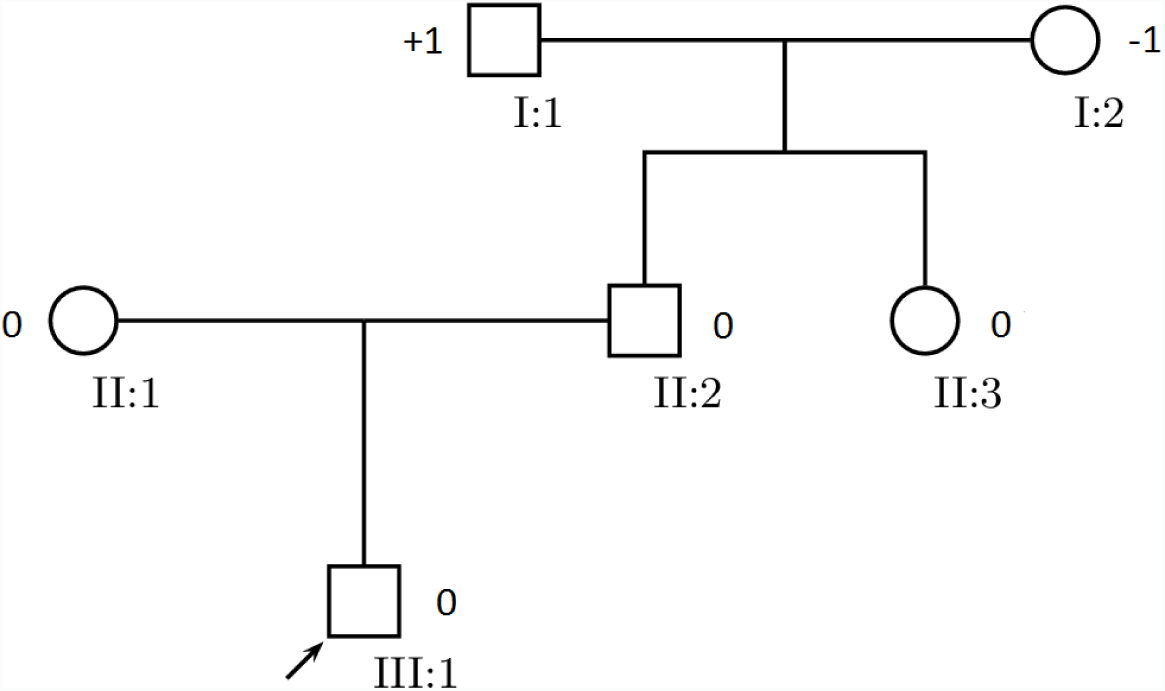
Assignment of bias to an Autosomal Recessive inheritance scheme.

Expected Bias from I Parents to II children: 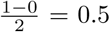

Bias from II children (II-2, II-3): (1-0.5) + (0-0.5) = 0

Expected Bias from II Parents to III children: 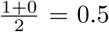

Bias from III children (III-1): (0-1.5) = −0.5^2^ = 0.25

Total Bias from all levels: 0 + 0.25 = 0.25

**Figure 9:**
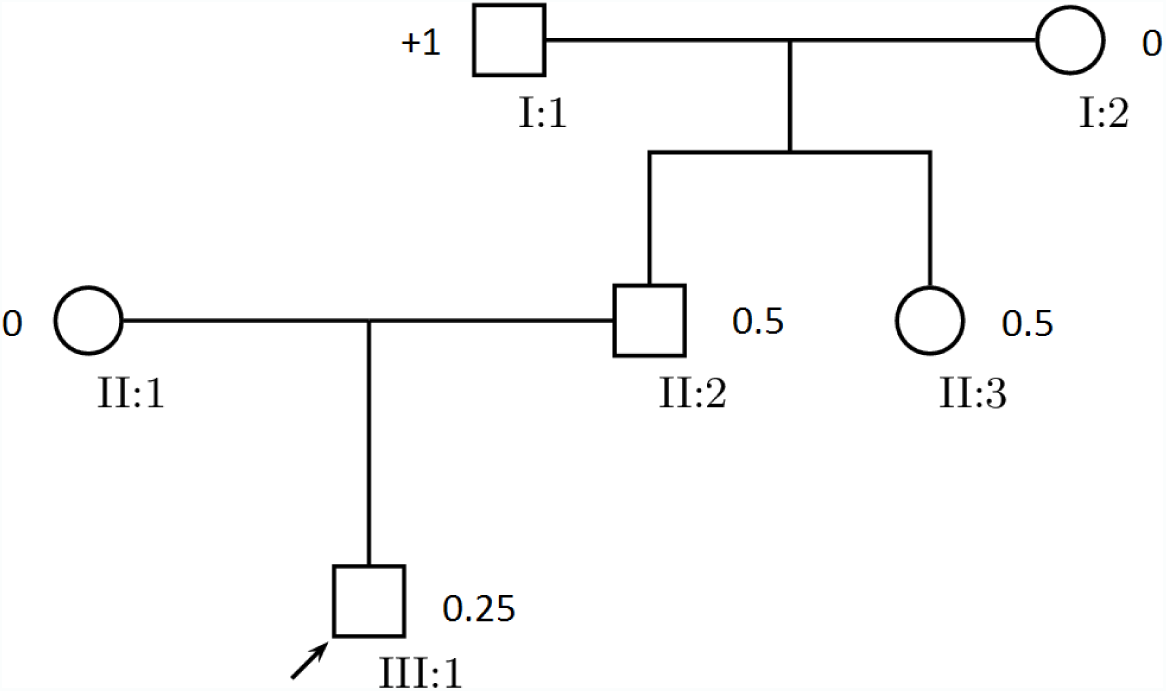
Assignment of bias to an Autosomal Dominant inheritance scheme.

Expected Bias from I Parents to II children: 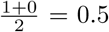

Bias from II children (II-2, II-3): (1-0.5) + (0-0.5) = 0

Expected Bias from II Parents to III children: 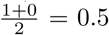

Bias from III children (III-1): (1-0.5) = −0.5^2^ = 0.25

Total Bias from all levels: 0 + 0.25 = 0.25

**Figure 10:**
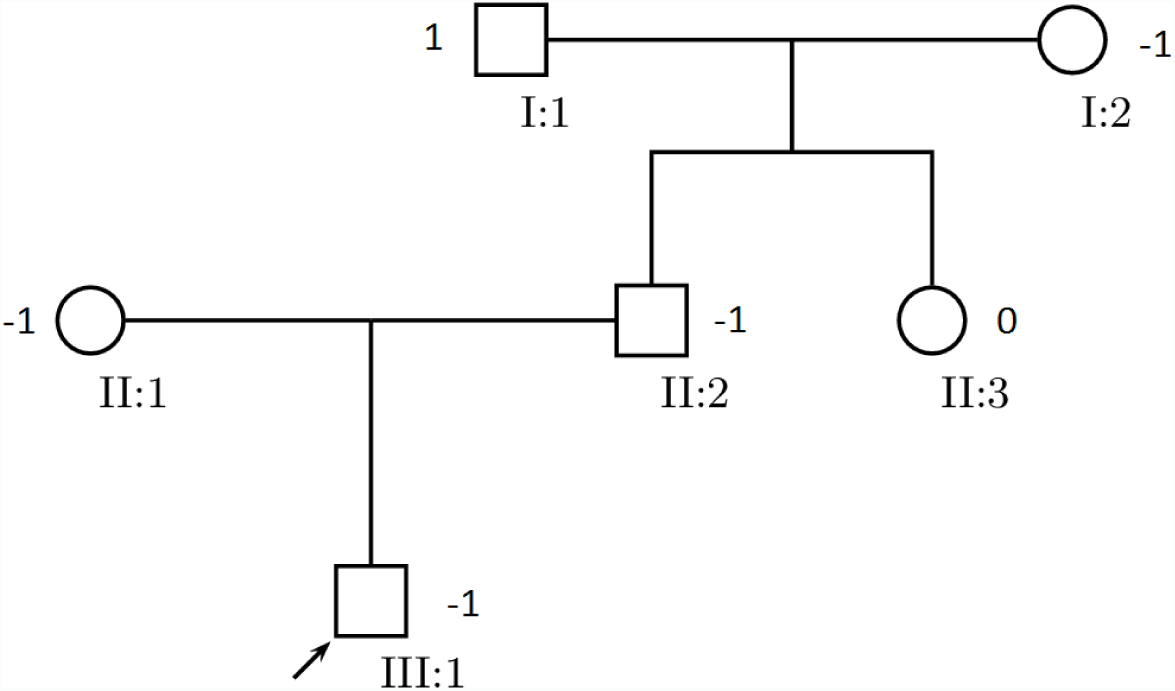
Assignment of bias to an X-Linked Recessive inheritance scheme.

Expected Bias from I Parents to II children: ♂ = 0, 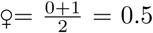

Bias from II children (II-2, II-3): (1-0) + (0-0) = 1^2^ = 1

Expected Bias from II Parents to III children: ♂ = 0, 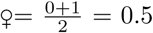

Bias from III children (III-1): (1-0) = 1^2^ = 1

Total Bias from all levels: 1 + 1 = 2

**Figure 11:**
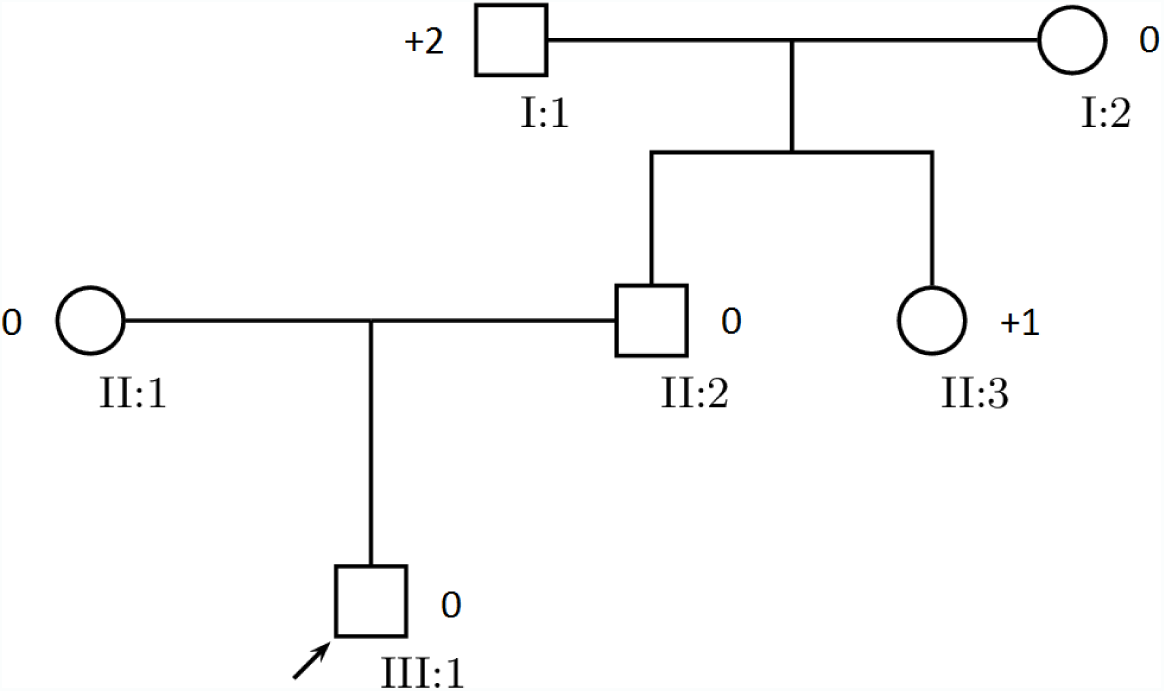
Assignment of bias to an X-Linked Dominant inheritance scheme.

Expected Bias from I Parents to II children: ♂ = 0, 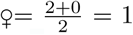

Bias from II children (II-2, II-3): (2-0)*1.5 + (0-1) = 3^2^ = 9

Expected Bias from II Parents to III children: ♂ = 0, 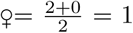

Bias from III children (III-1): (2-0)*1.5 + (0-1) = 3^2^ = 9

Total Bias from all levels: 9 + 9 = 18

**Figure 12:**
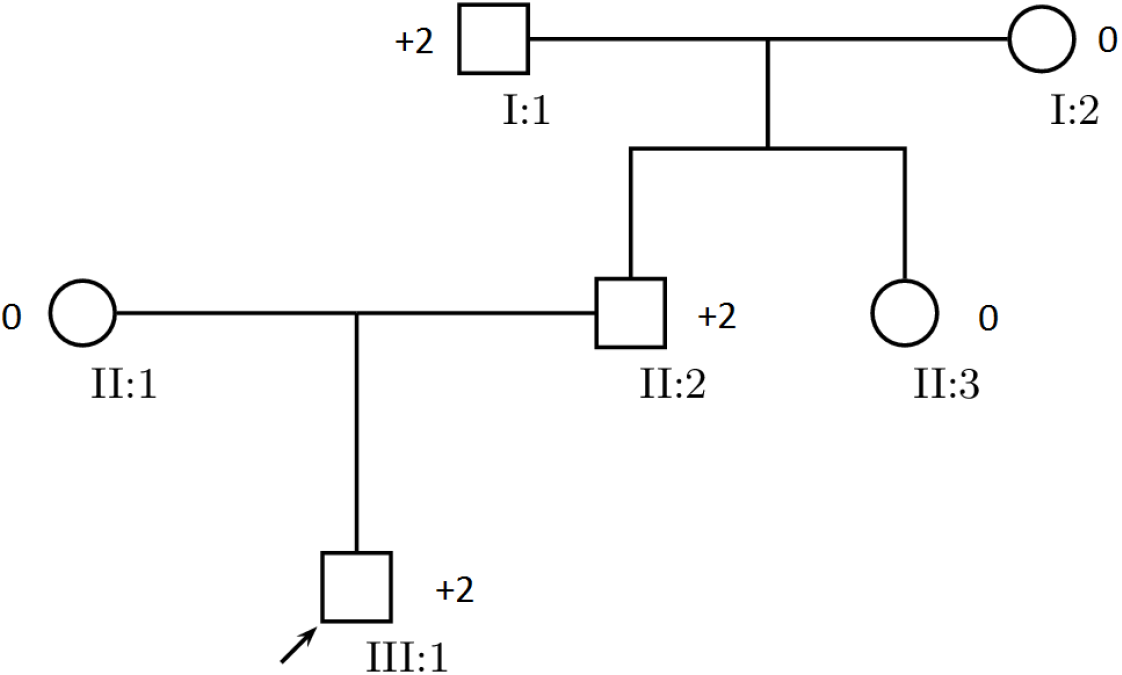
Assignment of bias to an Y-Linked inheritance scheme.

Expected Bias from I Parents to II children: ♂= 2, ♀= 0

Bias from II children (II-2, II-3): (2-2) + (0-0) = 0^2^ = 0

Expected Bias from II Parents to III children: ♂= 2, ♀= 0

Bias from III children (III-1): (2-2) = 0^2^ = 0

Total Bias from all levels: 0 + 0 = 0

After our calculations, we have the corresponding values for Autosomal Recessive, Autosomal Dominant, X-Linked Recessive, X-Linked Dominant, and Y-Linked: 6, 1.5, 12, 6, 0. Now, apply these values to a Chi-Square Distribution. Since we examined 2 parent-child systems (I-1/I-2 & II-1/II-2), our degrees of freedom is 2-1 = 1.

**Table 2:**
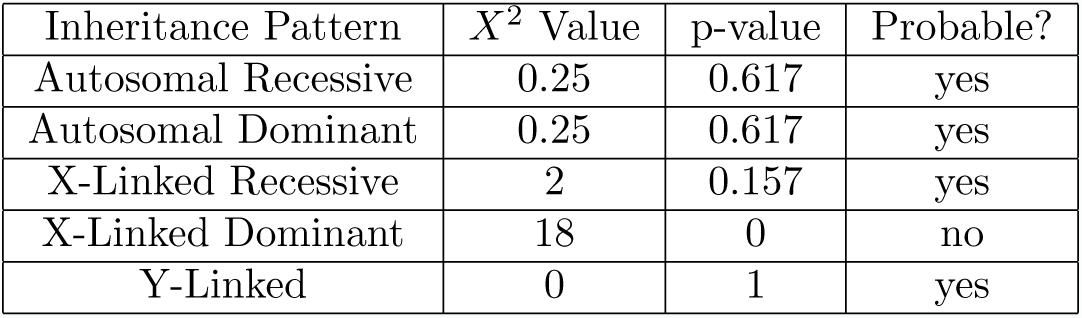
Analysis of Inheritance Patterns for Example 2.

Our results say Y-Linked, Autosomal Dominant, Autosomal Recessive and X-Linked Recessive inheritance systems are probable, with Y-Linked and Autosomal systems being much more likely. This pedigree is very ambiguous, as I have provided no information about heterozygosity. Though it may seem counter-intuitive, I did an ambiguous example to illustrate a point. When we generate models for this system, the self-correcting property allowed us to generate accurate models for Autosomal Systems in addition to the Y-Linked system. By allowing our algorithm to strategically assign heterozygotes, we were able to maximize the probability that a model might fit. For an unknown pedigree, this property is very important for generating a “Best Fit” model.

However, Y-Linked is more probable because the Autosomal and X-Linked systems depend on chance to achieve the pedigree above. For example, in the Autosomal Dominant inheritance scheme, it is possible for I-1 to pass it’s dominant allele to II-2, then II-2 to pass that dominant allele to III-1. However, this is much less likely since there is a chance that the male may inherit the unaffected allele from their mother. Since this must happen twice (I generation and II generation), those inheritance systems are less likely.

### Example 3 (X-Linked Recessive)

Finally, I will use a complicated example utilizing the history of Hemophilia in the royal family[6]. The data I am using is pulled from Hugo Iltis’ paper, “Hemophilia, ‘The Royal Disease’”. This data is inherently flawed. However, I seek to show that my model is still finds the best fit.

**Figure 13:**
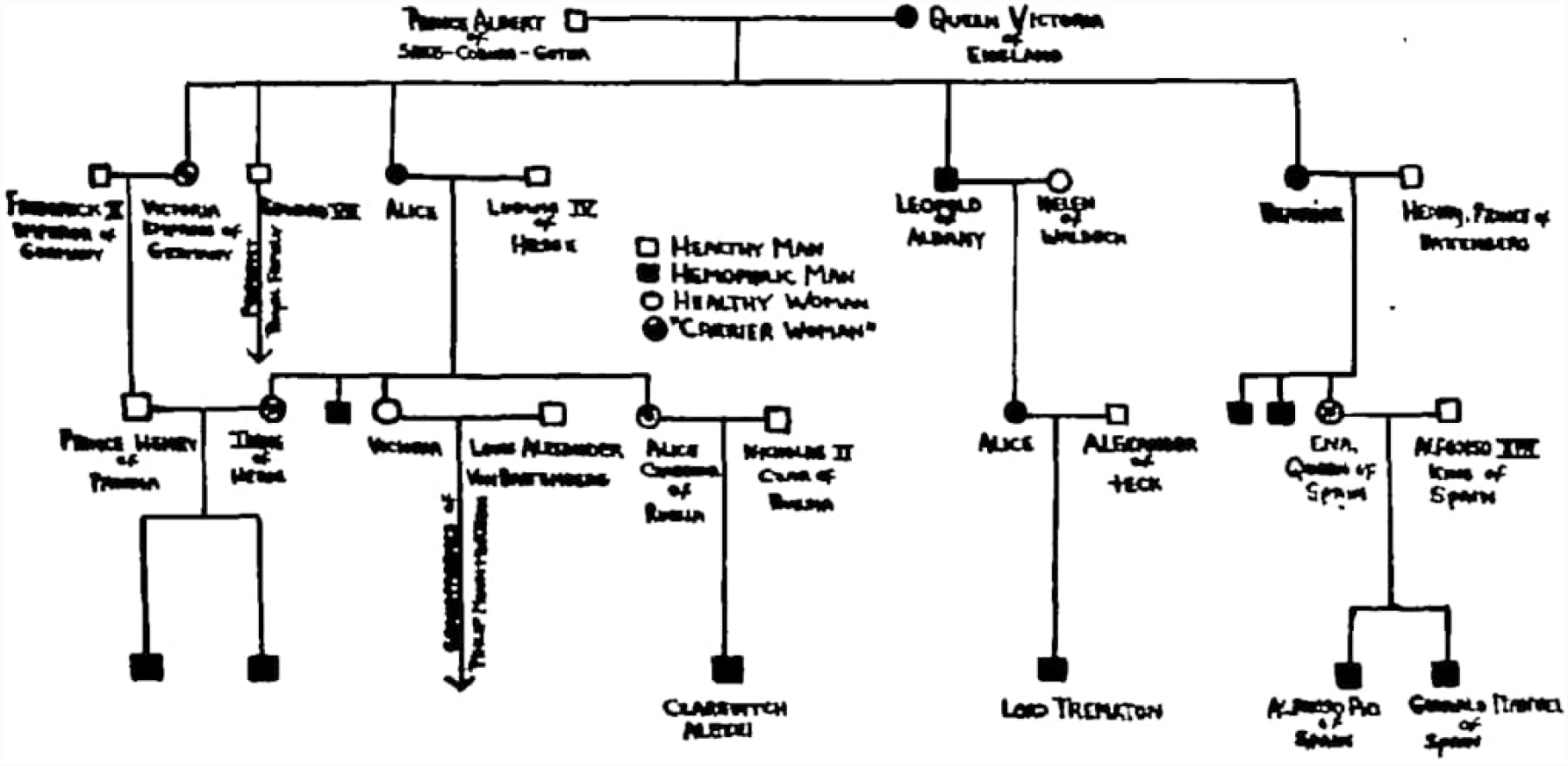
Sample Hemophilia pedigree taken from “Hemophilia, ‘The Royal Disease’”.

Let:

*α* be Autosomal Recessive

*β* be X-Linked Recessive

*γ* be Autosomal Dominant

*δ* be X-Linked Dominant

*ε* be Y-Linked

Δ to be *δ*_*k*_ *− ϕ*_*k*_, the difference between expected and observed for an individual.

Best for Individual - Smallest variance for individual.

Best So Far - Smallest total variance up to the individual.

**Table 3:**
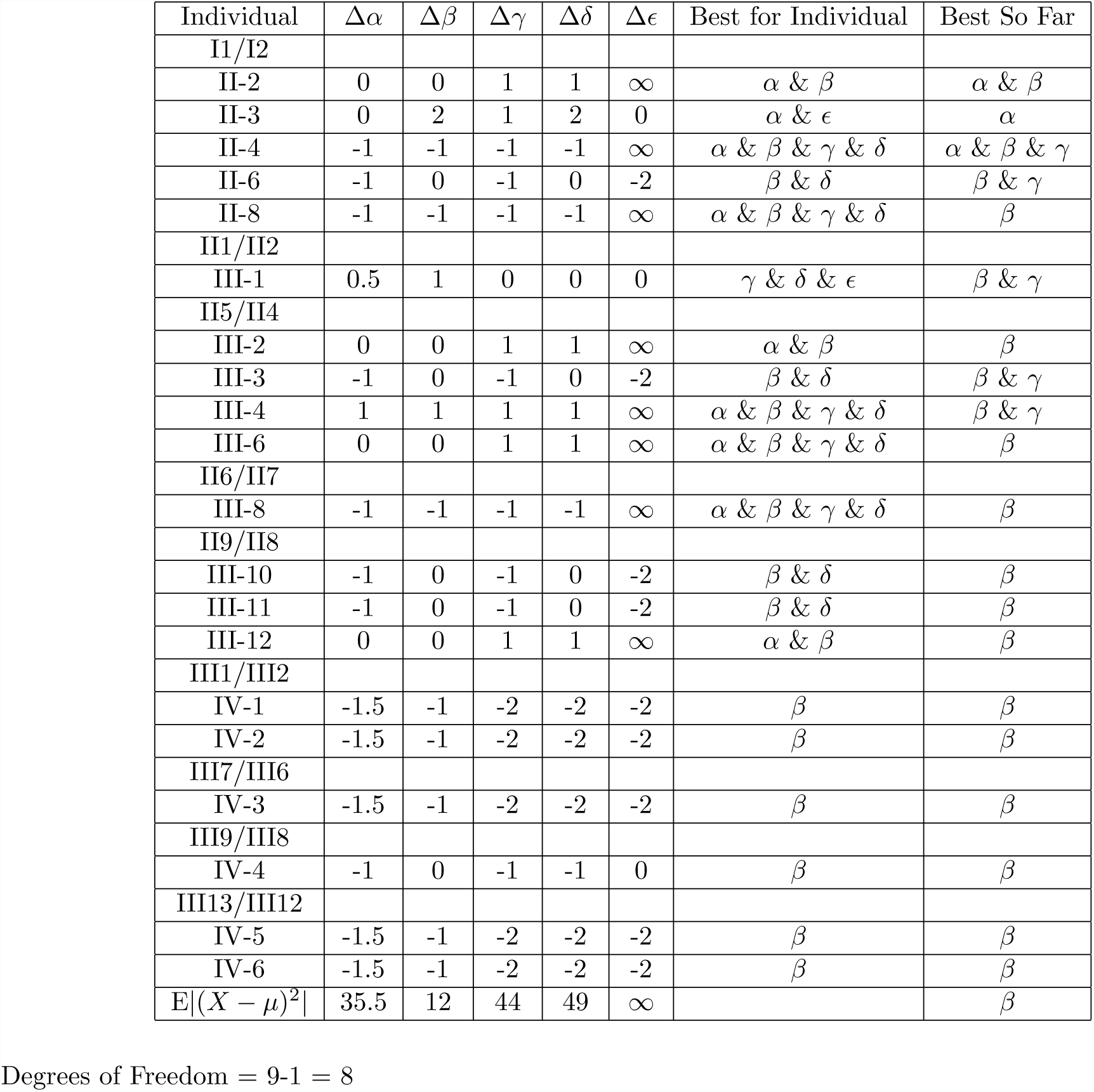
Scoring matrix for hemophilia.

**Table 4:** Analysis of Inheritance Patterns for Example 3.

| Inheritance Pattern | $X^2$ Value | p-value | Probable? |
| --- | --- | --- | --- |
| Autosomal Recessive | 35.5 | 0 | no |
| X-Linked Recessive | 12 | 0.151 | yes |
| Autosomal Dominant | 44 | 0 | no |
| X-Linked Dominant | 49 | 0 | no |
| Y-Linked | ∞ | 0 | no |

### Analysis

We can say with 15.1% certainty that this pedigree is X-Linked Recessive. Autosomal Recessive, Autosomal Dominant, X-Linked Dominant, and Y-Linked are not probable.

From our “Best so Far” column, we can track how each inheritance system rates as we add more data. Generation II has many problems. Since the inhertiance for Hemophilia is X-Linked Recessive, a cross between an affected female and unaffected male should produce all carrier females and all affected males. However, we can already see this break down in generation II. For example II-4 or II-8 are affected females which must have received two copies of the affected allele. [6] However, since the father is unaffected, he cannot possibly pass a affected allele to his daughters. Therefore, Autosomal Dominant becomes more probable for a brief moment. If the system were Autosomal Dominant, the mother has a chance of passing the affected allele to her daughters, causing them to be affected.

However, as we continue to add more data, it is clear that X-Linked Recessive is more accurate. Generation II does not seem to be X-Linked Recessive, and as such, Autosomal Dominant garnered the least variance. Though most of Generation II is ambiguous, the later generations favor X-Linked Recessive heavily. For example, the family produced from II-9 and II-9 fits X-Linked Recessive inheritance perfectly. This is reflected in our scoring matrix, as X-Linked Recessive is the only system to score zero variance for this family (III-10, III-11, III-12). Additionally, we can see that this algorithm was able to correctly identify generation IV to be X-Linked Recessive. The X-Linked Recessive system scored less variance than all other systems for Generation IV, therefore X-Linked Recessive is the most accurate system for Generation IV. This can be confirmed by a manual analysis. Many of the affected males in Generation IV have a carrier mother and an unaffected father (III-1/III-2, III-6/III-7, III-12/III-13). This does not fit Dominant inheritance schemes, as both parents are unaffected. An affected child cannot have two unaffected parents. These results also do not fit Autosomal Recessive inheritance. A carrier mother and unaffected father should always produce unaffected progeny, as the child will always inherit a working allele. Finally, the affected sons in Generation IV could not inherit an affected allele from an unaffected father, so Y-Linked is also inaccurate.

I have compiled and attached two matrices that represent expected and observed biases.

## Discussion

### Benefits

As a numerical system, this implementation offers many of the same benefits that others of the same category do. For example, finding the probability for individuals to be affected can be made much quicker than a standard pedigree because the calculation only need the biases of the parents, and, if self-correcting is needed, the biases of the siblings. This drastically improves running time in large pedigrees. For example, let us have a pedigree of depth D with N individuals in the last generation. Let search refer to finding the chance of an individual being affected, and insert be the probability of an individual having an affected child (genotype of mate is known).

**Table 5:** Run Time costs.

|  | Worst Case Search | Best Case Search | Worst Case Insert | Best Case Insert |
| --- | --- | --- | --- | --- |
| Standard Pedigree | $O(D)$ | $O(1)$ | $O(D)$ | $O(1)$ |
| Improved Pedigree | $O(1)$ | $O(1)$ | $O(1)$ | $O(1)$ |

This improved system also considers information that is not present but can be inferred. For example, suppose we have an Autosomal Recessive disease, with one grandparent who is affected and one who is not. Suppose that we know the mother of an individual is an outsider who mates with a son of the grandparents. However, we have no information about the father or others in his generation. Under this system, we would still know that the father is definitely a carrier, as his bias will be 0. Though this example may seem trivial, consider the case where we do not know generation 4 of a 16 generation pedigree. This implementation will still track the contribution of generation 4 to their descendants, which is incredibly useful.

Finally, this model allows us to calculate the probability that certain inheritance systems are present. This model is flexible, and allows us to know if there is a possibility of multiple inheritance systems instead of trying to pick one. It also accounts for ambiguity in heterozygotes and factors this into its calculations for inheritance systems. Accounting for ambiguity is incredibly useful if we are only given phenotypic data.

### Limitations

As I developed these algorithms theoretically, they have many foreseeable limitations.

First and foremost are the assumptions I make. There is no guarantee that outsiders will be what I assume them to be (Unaffected are Homozygous Dominant in Autosomal Recessive scheme, Affected are Heterozygous in Autosomal Dominant scheme, etc…). Every time that the assumption is wrong, the algorithm must back track and rectify the mistake (Self-Correcting property). Secondly, there is no distinction between having two Heterozygotes as parents versus a Homozygous Dominant and Homozygous Recessive. Though the allelic frequencies are equal, the two sets of parents have different effects. If a disease is Autosomal Recessive, and two carriers have a child, there is a 25% chance the child being affected. If a Homozygous Dominant and Homozygous Recessive couple have a child, it will always be unaffected. My system does not account for this. Additionally, my models do not account for Non-Mendelian traits, nor traits that are linked to other traits. Using these models to determine inheritance is also inefficient. I have not implemented a threshold to stop tracking inheritance systems, but that is a possible improvement. For example, if an affected female is present in our pedigree, Y-Linked inheritance will be ruled out and we will no longer keep track of that inheritance scheme.

## Acknowledgements

I would like to thank Nathaniel Johnson for his statistics distribution website. This was key for analyzing the variance I obtained from my calculations. I would also like to thank Boris Veytsman and Leila Akhmadeeva for their online pedigree tool. This tool was both useful and highly beautiful, and I’m not sure if I could have undertaken this project without it. Finally, I would like to thank David Dinata for helping me structure this paper.

